# DSResSol: A sequence-based solubility predictor created with dilated squeeze excitation residual networks

**DOI:** 10.1101/2021.08.09.455643

**Authors:** Mohammad Madani, Kaixiang Lin, Anna Tarakanova

## Abstract

Protein solubility is an important thermodynamic parameter critical for the characterization of a protein’s function, and a key determinant for the production yield of a protein in both the research setting and within industrial (e.g. pharmaceutical) applications. Thus, a highly accurate in silico bioinformatics tool for predicting protein solubility from protein sequence is sought. In this study, we developed a deep learning sequence-based solubility predictor, DSResSol, that takes advantage of the integration of squeeze excitation residual networks with dilated convolutional neural networks. The model captures the frequently occurring amino acid k-mers and their local and global interactions, and highlights the importance of identifying long-range interaction information between amino acid k-mers to achieve higher performance in comparison to existing deep learning-based models. DSResSol uses protein sequence as input, outperforming all available sequence-based solubility predictors by at least 5% in accuracy when the performance is evaluated by two different independent test sets. Compared to existing predictors, DSResSol not only reduces prediction bias for insoluble proteins, but also predicts soluble proteins within the test sets with an accuracy that is at least 13% higher. We derive the key amino acids, dipeptides, and tripeptides contributing to protein solubility, identifying glutamic acid and serine as critical amino acids for protein solubility prediction. Overall, DSResSol can be used for fast, reliable, and inexpensive prediction of a protein’s solubility to guide experimental design.

**Availability:** The source code, datasets, and web server for this model are available at https://github.com/mahan-fcb/DSResSol

## 1. Introduction

Solubility is a fundamental protein property, which can give useful insights into the protein’s function or potential usability, for example, in foams, emulsions and gels [1] and therapeutics applications such as drug delivery [2]. In practice, the analysis of protein solubility is the most important determinant of success (i.e. high yields) in therapeutic protein and protein-based drug production[3], [4] In the research setting, producing soluble recombinant protein is essential for investigating the functional and structural properties of the molecule [5]. To improve yields experimentally, there exist certain refolding methods that utilize weak promoters and fusion proteins or optimize expression conditions, e.g. by using low temperatures [3], [4]. However, these methods cannot ensure production of soluble protein from a relatively small trial batch size limited by production cost and time. Given these concerns, reliable computational procedures for discovering potentially soluble protein targets for experimental testing can help to avoid expensive experimental trial and error approaches.

A protein’s structure and sequence features such as the isoelectric point, polarity, hydrophobicity, turn-forming amino acids, etc., are crucial intrinsic factors in protein solubility determination [6]–[8]. On this basis, several in silico approaches have been developed to predict protein solubility by using the protein sequence and its features. The majority of these tools use traditional machine learning models such as support vector machines (SVM) [9] and gradient boosting machines [10], employing pre-extracted features (i.e., features that are extracted from the protein sequences via other bioinformatics tools before feeding them into machine learning models) as input for these models. For example, SOLpro employs two-stage SVM models for training 23 extracted features from the protein sequences [4]. PROSO II utilizes a two-layered structure, including Parzen window [11] and first level logistic regression models as the first layer and a second-level logistic regression model as the second layer [12]. In more recent models such as PaRSnIP [13] gradient boosting machine models are used. This predictor utilized the frequency of mono-, di-, and tripeptides from the protein sequence in addition to other biological features such as secondary structure and fraction of exposed residues in different solvent accessibility cutoffs, as training features. SoluProt is the newest solubility predictor using a gradient boosting machine for training [14]. To evaluate the performance of this tool, a new independent test set was utilized. Notable, the frequency of important dimers extracted from the protein sequences was used as input features of SoluProt model [14]. All of the aforementioned models are two stage models, with a first stage set up for extracting and selecting features and a second stage employed for classification. Deep learning (DL) models circumvent the need for a two-stage model. DeepSol is the first deep learning-based solubility predictor proposed by Khurana and coworkers [15] built as a single stage predictor through the use of parallel convolutional neural network layers [16] with different filter sizes to extract high dimensional structures encoding frequent amino acid k-mers and their non-linear local interactions from the protein sequence as distinguishable features for protein solubility classification [15].

In this study, we propose a novel deep learning framework to create a sequence-based solubility predictor that outperforms all currently available state-of-the-art predictors: DSResSol (Dilated Squeeze excitation Residual network Solubility predictor). The main difference between our model and DeepSol is in the utilization of parallel squeeze excitation residual networks with a specific architecture, resulting in a significant improvement in performance of DSResSol compared to DeepSol. In contrast to DeepSol, our framework can easily capture key global interactions between amino acid k-mers through this novel architecture. Specifically, we employ parallel Squeeze-and-Excitation residual network blocks that include dilated convolutional neural network layers (D-CNNs) [17], residual networks blocks (ResNet) [18] and Squeeze-and-Excitation (SE) neural network blocks [19] to capture not only extracted high dimensional amino acid k-mers from the input protein sequence but also both local and long-range interactions between amino acid k-mers, thereby increasing the information extracted by the model from the protein sequence. Our work is inspired by recent studies using dilated convolutional neural networks and SE-ResNet for protein sequence and text classification [20], [21].

The traditional method to capture long-range interactions in data and to solve vanishing gradient problems is to use Bidirectional Long Short-Term (BLSTM) memory networks [22], [23]. However, BLSTM implementation significantly increases the number of parameters in the model. Thus, we use D-CNNs instead of BLSTM because D-CNNs perform as simple CNN operations but over every *n^th^* element in the protein sequence, resulting in captured long-range interactions between amino acid k-mers. ResNet is a recent advance in neural networks that utilizes skip connections to jump over network layers to avoid the vanishing gradient problem, and gradually learns the feature vectors with many fewer parameters [18]. The Squeeze-and-Excitation (SE) neural network blocks explicitly model interdependencies between channels thereby directing higher importance to specific channels within the feature maps, over others. Thus, by designing a novel architecture that combines these three neural networks in a specific manner together with parallel CNNs layers, we build a highly accurate solubility predictor. In the first model instance, DSResSol (1), we use only protein sequence as input and protein solubility as output. In the second model instance, DSResSol (2), we include pre-extracted biological features added to the model as a hidden layer to improve the model’s performance. We further include an analysis of the most important single amino acids, dipeptides, and tripeptides for protein solubility based on model results, and compare with experimental findings.

## 2. Materials and Methods

### 2.1 Data preparation and feature engineering

To create the training set, we use the Target Track database [24]. Based on the methods proposed in previous studies [4], [12], we infer the solubility value for each protein sequence within the training set. A protein is labeled as insoluble if it cannot be expressed or purified experimentally. On the other hand, a protein is considered as soluble if it is realized as soluble, purified, crystallized, etc., e.g. an experimental state requiring the protein to be soluble. To maintain the generality of our training set, in contrast to previous studies such as SoluProt [14], we do not impose a limitation on the expression system for selecting the proteins included in our training set. To reduce the noise and redundancy from our training set, the following tasks are performed: (1) removing the transmembrane proteins based on the annotations from the Target Track database; (2) removing the proteins considered as insoluble but associated with a PDB structure; (3) eliminating protein sequences from the training set with a sequence identity of more than 25% via CD-HIT [25] to avoid any bias because of homologous sequences within the training and testing sets. Finally, we use a fairly balanced training set that includes approximately the same number of proteins within the soluble and insoluble classes. Thus, in total, our final training set contains 40317 protein sequences including 19718 soluble and 20599 insoluble protein sequences.

For model evaluation, we utilize 2 different independent test sets. Both test sets include proteins that have been expressed in E. coli:

1. The first test dataset was proposed by Chang et al. [26]. This dataset includes 2001 protein sequences and their corresponding solubility values.
2. The second test dataset, first proposed by Hon et al. [14], has been constructed from a dataset generated by the North East Structural Consortium (NESG) [27], and includes 9644 proteins expressed in E. coli. The original dataset consists of integer values in the range of 0 to 5 for levels of expression and soluble fraction recovery. We maintain consistency between the procedure for constructing the training set and test set. Finally, similar to the SoluProt test set [14], we transform the solubility level of each protein within the NESG test set to a binary value.

To decrease the overlap between sequences within both test sets and the training set, all protein sequences in the training set which have sequence identity of more than 15% with protein sequences in both test sets are eliminated. This significantly reduces redundant sequences in our training set. SI Table 1 catalogues the dataset construction/reduction process for both the training set and both test sets in detail.

### 2.2. Model architecture

Protein solubility prediction is a binary classification problem. Within the datasets, each protein sequence is assigned a solubility value equal to 0 or 1. Thus, the solubility propensity for each sequence evaluated by our model is assigned a score in the range [0,1]. The DSResSol (1) model includes 5 architectural units, including a single embedding layer, nine parallel initial CNNs with different filter sizes, nine parallel SE-ResNet blocks, three parallel CNNs, and fully connected layers, sequentially (Figure 1). The architecture of DSResSol (2) is equivalent to DSResSol (1). DSResSol (2) has an additional input layer to receive 85 additional biological features.

**Figure 1.**
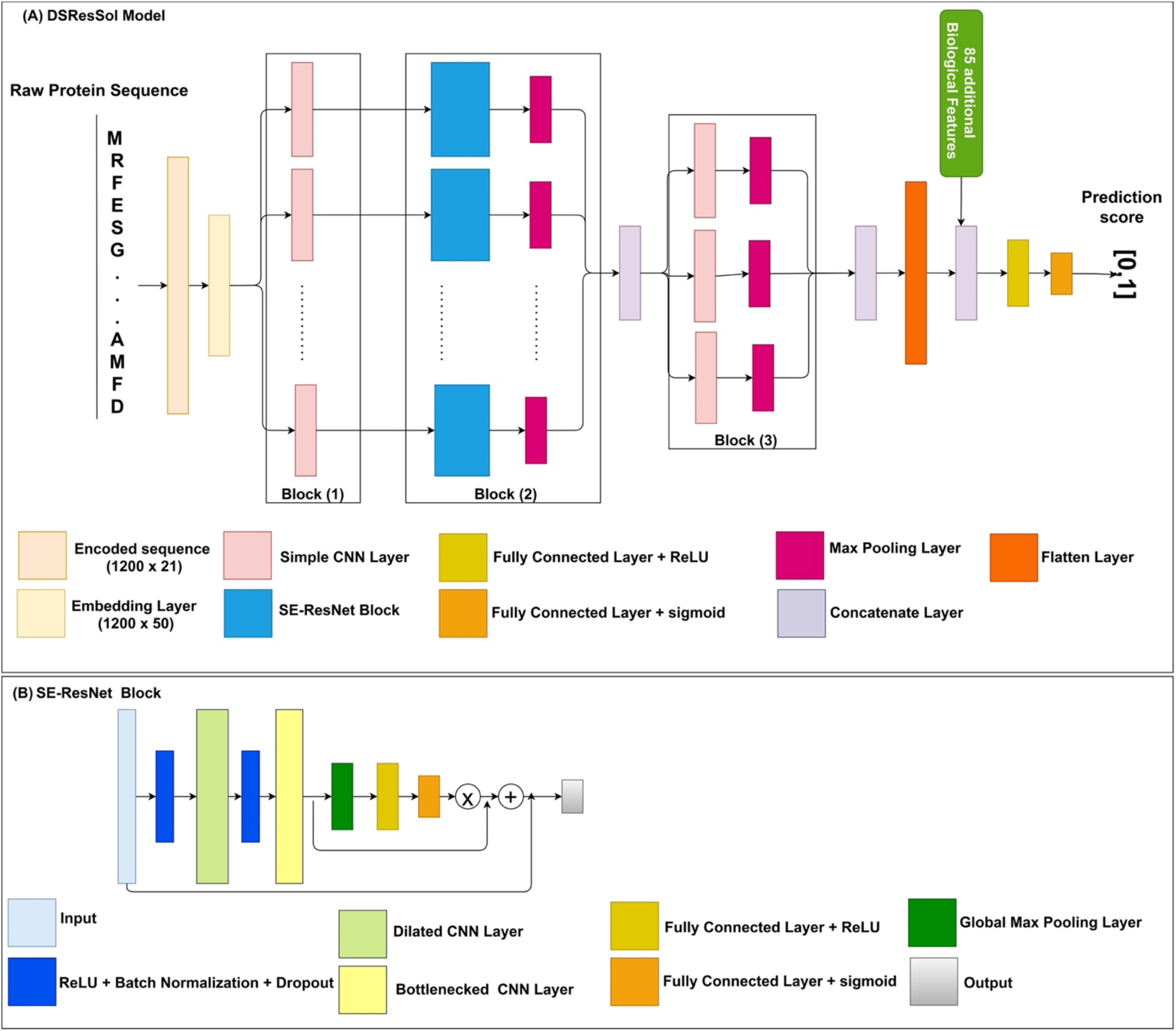
Schematic of the DSResSol model architecture. (A) DSResSol Block 1 has 9 initial CNNs with 32 filters within each CNN and filter sizes from k = 1 to 9; Block 2 includes 9 SE-ResNet blocks; Block 3 has another 3 CNNs with 32 filters with filter sizes {11, 13, 15ļ. 85 additional biological features are used in DSResSol (2) only. (B) Schematic of the SE-ResNet block architecture. Each SE-ResNet block has a single dilated CNN with dilation rate of 2 and filter size of 3, and single bottlenecked CNN with filter size of 1. All Max Pooling layers have 1 stride and window size of 3.

#### 2.2.1 Model input (encoding protein sequences)

To use protein sequences as inputs of the DSResSol model, we employ two major preprocessing approaches. First, protein sequences are parameterized to the vectors *X* = {*x*_0_, *x*_1_, *x*_2_,…,*x_L_*} where *x_i_* ∈ {0,1,2,, 20}. The numbers from 1 to 20 represent amino acid residues, and 0 indicates a gap. Second, each sequence is padded to the fixed-length vector having length L=1200 to generate same-sized vectors.

#### 2.2.2 Embedding layer

This layer is defined as a lookup table for mapping and transforming the discrete input into the continuous fixed-sized vectors, so-called embedding vectors. Thus, during the training process, a continuous feature is learned from each amino acid. The embedding layer transforms input sequence vector x∈ R^(1200×21) to a dense continuous feature representation via embedding weight matrix W_e ∈ R^(50×21). The output of embedding layer, i.e., the feature map, is E=x×W_e. The embedding dimension is 50. Note that training the W_e happens along the whole network.

#### 2.2.3 Nine Parallel CNNs

The feature map is fed to nine CNN layers with different filter sizes k from 1 to 9: k=[1,2,…,9} (Figure 1 (A) Block (1)). We use different filter sizes in the CNNs to extract amino acid k-mers, i.e., “biological words,” with different sizes between one (monopeptide) to nine (nonapeptide) from the input sequences. This component of the model is inspired by DeepSol [15]. In fact, the filter size in the CNN is equal to the convolutional window size along the characters of the sequence.

#### 2.2.4 SE-ResNet blocks with dilated CNNs

A Squeeze-and-Excitation residual network (SE-ResNet) block consists of two main parts: a residual network part and a Squeeze-and-Excitation block, linked via a residual connection (Figure 1 (A) Block (2)).

##### 2.2.4.1 ResNet

Residual neural networks (ResNet) [18] are an outstanding new discovery for neural networks making neural networks deeper, by resolving the gradient vanishing problem with fewer parameters than traditional neural networks such as CNNs. In ResNet, the gradients can flow directly through the skip connections backwards from later layers to initial filters [18], [29]. ResNets are utilized in a wide range of applications including natural language processing [30] and image classification [18].

The architecture of ResNet in the DSResSol model contains two CNNs, a dilated CNN [31] and a bottlenecked CNN [21], followed by batch normalization and a rectified linear unit as the activation function [32]. Figure 1 (B) shows the ResNet block within the SE-ResNet module. Using a bottlenecked convolution layer speeds up the computation and increases the ResNet block’s depth by using fewer parameters and a thinner ResNet block [33]. An n-dilated convolution captures local and global information about amino acid k-mers without significantly increasing the model parameters because it behaves like a simple convolution operation over every n^th element in a sequence [21]. In fact, this type of convolution enables the model to capture long-range interactions across the sequence. The dilated convolution utilizes kernels, which have holes. In this way, not only is the overall receptive field of convolution wider, but also the complexity and number of parameters are reduced [17], [21]. The utilization of n-dilated convolution is inspired by a recent study for protein family classification from protein sequence [21].

##### 2.2.4.2 SE Block

In addition to ResNet, each SE-ResNet block consists of a single Squeeze-and-Excitation (SE) block. The SE block focuses on more important channels within the feature maps. In other words, the Squeeze-and-Excitation blocks can recalibrate the channels in the learned feature maps, which results in stimulating more important channels and hindering weak channels within the feature maps [34], [35]. The feature input of the SE block is passed through the Global Max Pooling layer. This layer reduces each channel in the input feature map to a single value, which is the maximum value within each channel. Suppose the input tensor of the SE block has the shape of L×C, where C is the number of channels within the feature map and L is the feature dimension. After passing this tensor through the Global Max Pooling operator, the shape of the output will be reduced to C×1. To map adaptive scaling weights for the output of the Global Max Pooling layer, we employ two fully connected (FC) layers. In the first FC layer, the number of units is set to C/8, and the activation function is the rectified linear unit (ReLU) [32]. In the second FC layer, the number of units is set to C to project back the first FC layer’s output to the same dimensional space as the input, returning to C neurons. In summary, the C×1 tensor input is passed through the first FC layer; next, a weighted tensor of the same shape is obtained from the second FC layer as output. The sigmoid is utilized as the second FC activation function to scale the value to a range of 0 to 1. Using a simple broadcasted element-wise multiplication, the second FC layer’s output is applied to the SE block’s initial input [34]. To complete the SE-ResNet block, the SE block’s output is concatenated with the ResNet block’s input (Figure 1 (A)). These processes are the same for each SE-ResNet block. The feature maps derived from each SE-ResNet block are fed to the Max Pooling layer to accumulate feature maps by taking maximum values over the sub-region along with the feature map. The output Max Pooling layers are merged to generate feature maps for the next layer.

#### 2.2.5 Last three CNNs

For the next stage of the model, the three convolution layers with a filter size of 11, 13, and 15, respectively, following three Max Pooling layers (Figure 1 (A) Block (3)). This stage is responsible for extracting more contextual features from the merged outputs of the SE-ResNet module. Finally, all three feature maps obtained from this stage are concatenated.

#### 2.2.6 Fully connected layers

The output of the previous stage is flattened to a 1D array then fed into a single FC layer with hidden neurons of size 128 and ReLU as the activation function. The final FC layer with sigmoid as the activation function generates the probability score for solubility propensity.

#### 2.2.7 Additional features

Similar to PaRSnIP [13], we use secondary structure, solvent accessibility, structural order/disorder, and global sequence features as additional biological features. We obtain secondary structure (SS) and relative solvent accessibility (RSA) features through the SCRATCH webserver [36], order/disorder (O/D) information from the ESpritz webserver [37], and global sequence features from the python package modLAMP [38]. To calculate features from the SS and RSA, we employ the PaRSnIP procedure [13]. We calculate the fraction content of SS for 3- and 8-state SS. In addition, we obtain the fraction of exposed residues at different cutoffs (FER-RSA) from 0% to 95% with 5% intervals. The FER-RSA values are multiplied by the hydrophobicity of exposed residues to extract another RSA-based feature group. Unlike the PaRSnIP model, we introduce additional O/D-related features to our biological features set. For O/D-related features, we calculate the number of disordered regions limited to fewer than 5 amino acids, sized between 5 and 10 amino acids, and sized larger than 10 amino acids, as well as the frequency of each amino acid in disordered regions. In total, 85 biological features are extracted from the protein sequence. A summary of all biological features used in our model is presented in Figure 2. These 85 pre-extracted biological features are concatenated to the feature maps derived from the flattening layer before fully connected layers in DSResSol (2).

**Figure 2.**
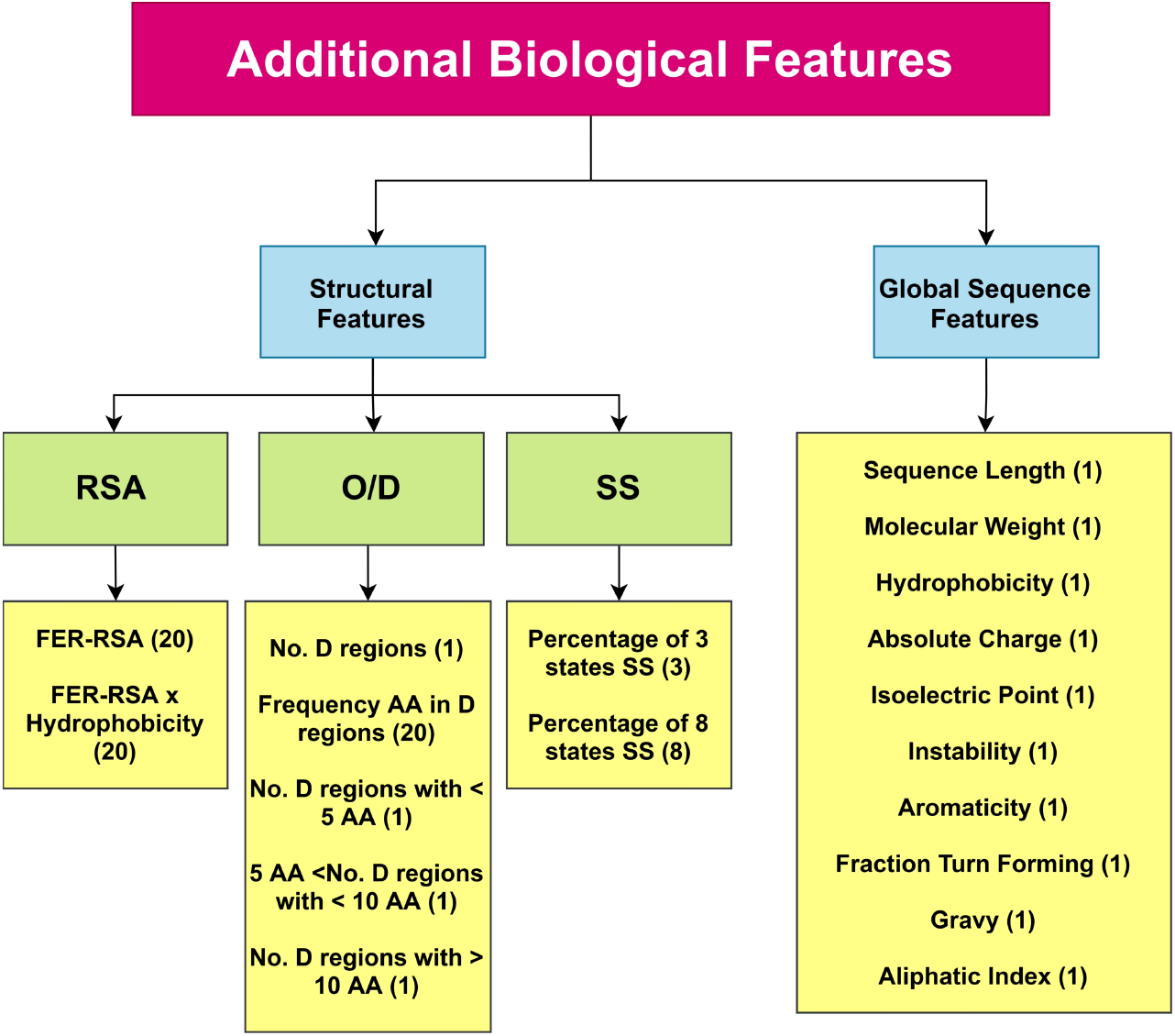
85 additional pre-extracted biological features are captured from the protein sequence and used for training the DSResSol (2) model. RSA: Relative Solvent Accessibility, SS: Secondary Structure, O/D: Order/ Disorder, FER-RSA: Fraction Exposed Residue at Relative Solvent Accessibility Cutoffs. The number of features for each type is shown in parentheses.

## 3. Training and Hyperparameter Tuning

The DSResSol model utilizes a binary cross-entropy [39] objective function to classify the protein sequence into two classes. Both models DSResSol (1) and (2) are fit for a different number of training epochs with the Adam optimizer [40]. Performance of the models depends on different hyperparameters such as: learning rate; number of; batch; size and number of filters in each convolution layer; embedding dimension; number of units in FC layers; etc. We have performed hyperparameter tuning by employing a grid search on 10-fold cross-validation similar to DeepSol [15]. SI Table 2 represents the tuned hyperparameters for both DSResSol (1) and DSResSol (2) models. After identifying optimal hyperparameters, we perform 10-fold cross-validation to train the models. We also use the early stopping approach during the training to avoid overfitting [15].

## 4. Evaluation metrics

To evaluate the performance of the DSResSol predictor in comparison with other predictors, we use accuracy, Matthew’s correlation coefficient (MCC), sensitivity, selectivity, and gain metrics as described in PaRSnIP [13].

## 5. Results

### 5.1 Model Performance

A 10-fold cross-validation is performed for the training process. In each cross-validation step, the training set is divided into ten parts. Nine parts are used for training and one part is used for validation. The performance of the DSResSol model is reported by using the ten models. To evaluate the stability in performance results, we use four different metrics: accuracy, precision, recall, and F1-score. Figure 3 represents the box plots of these metrics for all ten models obtained through 10-fold cross-validation for both DSResSol (1) and DSResSol (2) on both independent test sets. Notably, the variance in box plots corresponding to each metric for both DSResSol (1) and DSResSol (2) is very small, highlighting the outstanding stability in performance of the DSResSol predictor. For example, for DSResSol (2), among 10 models, the best model has accuracy 79.2% and weakest model has accuracy 78.4%, with a variance of 0.8% (SI Table 3 and 4).

**Figure 3.**
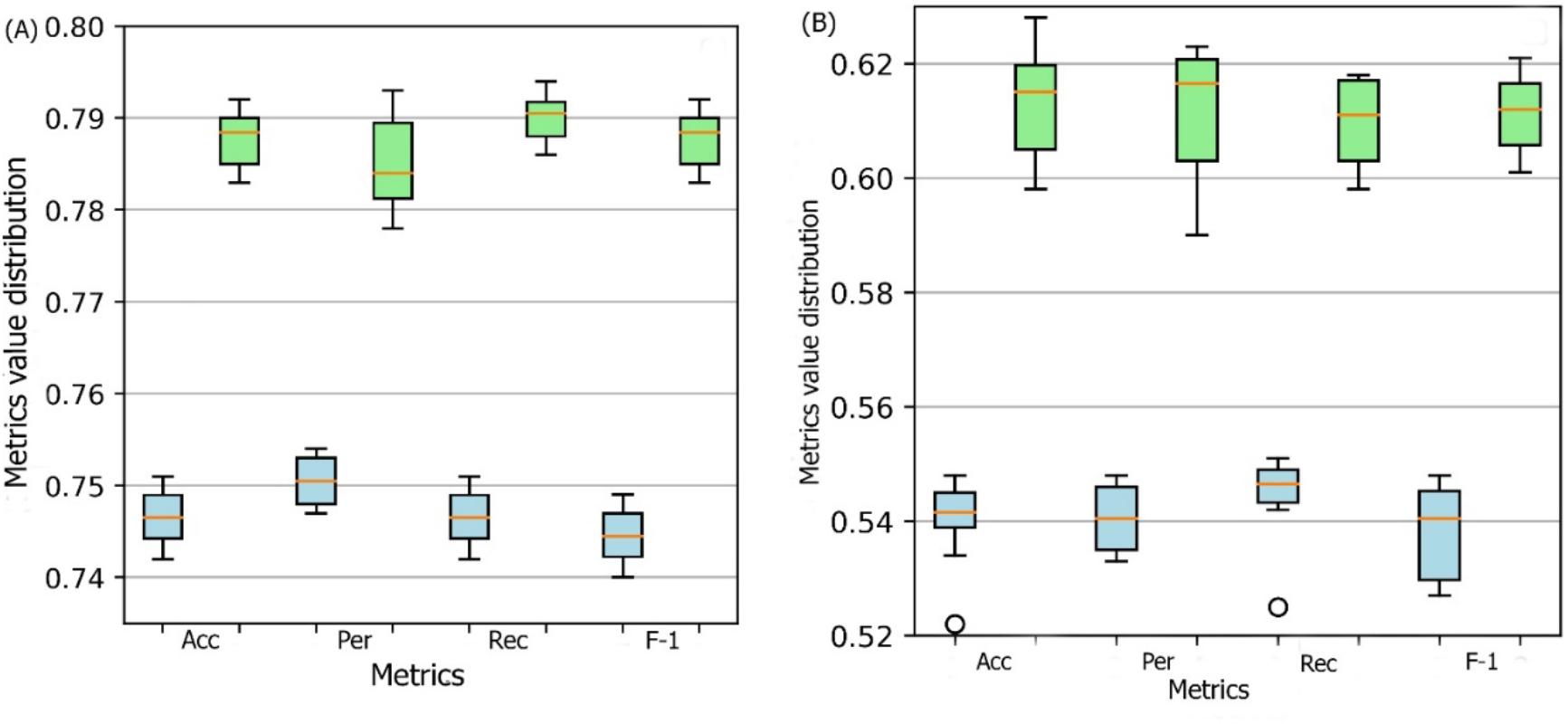
Box plot for 10 models obtained from 10-fold cross-validation for both DSResSol (1) and DSResSol (2) considering four metrics: ACC (accuracy), Per (precision), Rec (recall), and F-1 (f-1 score) for (A) Chang et al. test set [41], (B) NESG test set. Note: blue and green box plots represent the score distribution for DSResSol (1) and DSResSol (2), respectively.

To compare the performance of DSResSol with the best existing prediction tools, we use two different testing sets, the first one proposed by Chang et al. [41] and the second one, NESG dataset, proposed by Price et al [27] and refined by Hon et al. [14]. Table 1 and 2 display the performance of eight solubility predictors on both test sets. DSResSol (2) outperforms all available sequence-based predictor tools when the performance is assessed by accuracy, MCC, sensitivity for soluble proteins, selectivity for insoluble proteins, and gain for insoluble proteins.

**Table 1.** Performance of DSResSol in comparison with known existing models on first independent test set [41]. Note: Best per-forming method in bold.

| Model | ACC | MMC | Selectivity (Soluble) | Selectivity (Insoluble) | Sensitivity (Soluble) | Sensitivity (Insoluble) | Gain (Soluble) | Gain (Insoluble) |
| --- | --- | --- | --- | --- | --- | --- | --- | --- |
| DSResSol (2) | **0.796** | **0.589** | 0.817 | **0.782** | **0.769** | 0.823 | **1.634** | **1.564** |
| DSResSol (1) | 0.751 | 0.508 | 0.786 | 0.722 | 0.691 | 0.813 | 1.572 | 1.445 |
| SoluProt | 0.682 | 0.382 | 0.701 | 0.670 | 0.722 | 0.643 | 1.403 | 1.342 |
| DeepSol S2 | 0.762 | 0.546 | **0.821** | 0.721 | 0.681 | **0.843** | 1.642 | 1.442 |
| DeepSol S3 | 0.760 | 0.543 | 0.801 | 0.725 | 0.707 | 0.822 | 1.602 | 1.451 |
| PaRSnIP | 0.720 | 0.472 | 0.761 | 0.723 | 0.698 | 0.743 | 1.522 | 1.446 |
| DeepSol S1 | 0.720 | 0.471 | 0.752 | 0.706 | 0.691 | 0.749 | 1.504 | 1.412 |
| PROSSO II | 0.638 | 0.345 | 0.671 | 0.682 | 0.693 | 0.662 | 1.342 | 1.365 |
| SCM | 0.600 | 0.214 | 0.650 | 0.572 | 0.422 | 0.773 | 1.301 | 1.145 |
| PROSO | 0.581 | 0.161 | 0.582 | 0.575 | 0.541 | 0.622 | 1.164 | 1.151 |
| CCSOL | 0.543 | 0.083 | 0.543 | 0.539 | 0.514 | 0.572 | 1.087 | 1.081 |
| RPSP | 0.520 | 0.032 | 0.522 | 0.517 | 0.447 | 0.588 | 1.044 | 1.035 |

**Table 2.** Performance of DSResSol in comparison with known existing models on NESG test set [14]. Note: Best performing method in bold.

| Method | ACC | MCC | Selectivity (Soluble) | Selectivity (Insoluble) | Sensitivity (Soluble) | Sensitivity (Insoluble) | Gain (Soluble) | Gain (Insoluble) |
| --- | --- | --- | --- | --- | --- | --- | --- | --- |
| DSResSol (2) | **0.629** | **0.273** | 0.606 | **0.663** | **0.73** | 0.53 | 1.212 | **1.326** |
| DSResSol (1) | 0.557 | 0.169 | 0.558 | 0.555 | 0.54 | 0.58 | 1.117 | 1.110 |
| SoluProt | 0.578 | 0.189 | 0.575 | 0.581 | 0.60 | 0.56 | 1.150 | 1.162 |
| PROSSO II | 0.565 | 0.143 | 0.578 | 0.555 | 0.48 | 0.66 | 1.157 | 1.110 |
| SWI | 0.558 | 0.142 | 0.545 | 0.580 | 0.70 | 0.42 | 1.090 | 1.160 |
| CamSol | 0.535 | 0.115 | 0.548 | 0.527 | 0.39 | 0.68 | 1.097 | 1.054 |
| ESPRESSO | 0.493 | 0.093 | 0.493 | 0.492 | 0.55 | 0.44 | 0.987 | 0.984 |
| rWH | 0.519 | 0.133 | 0.532 | 0.513 | 0.31 | 0.73 | 1.065 | 1.026 |
| DeepSol S2 | 0.546 | 0.132 | **0.894** | 0.560 | 0.22 | **0.88** | **1.788** | 1.120 |
| SOLpro | 0.500 | 0.089 | 0.500 | 0.500 | 0.48 | 0.52 | 1.000 | 1.000 |

It is worth noting that the reason for significant differences between performance of model on the first and second test sets is not due to overfitting or overtraining. The difference between accuracy in training and validation for both DSResSol 1 and DSResSol 2 is less than 1.5% (SI Table 3 and 4), suggesting that neither model has an overfitting problem. Therefore, we conclude that the difference between the performance in different test sets originates from the nature of the test sets. Specifically, the protein sequences within the second test set are expressed in E. coli. while, for training process, we used a training set that includes mixed proteins (expressed in E. coli or other host cells) to achieve a more comprehensive model (SI Table 1).

For the first test set, we find that only the sensitivity value for the insoluble class, and the selectivity value for the soluble class were slightly inferior to the close competitor, DeepSol S2 [15]. The accuracy and MCC of DSResSol (2) is higher compared to DeepSol by at least 4% and 7%, respectively (Table 1). In addition, the sensitivity of DSResSol (2) for both soluble and insoluble proteins is close in value, suggesting that the DSResSol (2) model can predict both soluble and insoluble protein sequences with high accuracy and minimal bias. This consistency is missing in DeepSol S2 and DeepSol S3, the most accurate predictors to date. Sensitivity of DSResSol (2) for insoluble protein is about 83% which is comparable to the current best predictor (DeepSol S2 = 85%). On the other hand, DSResSol (2) can identify soluble protein with higher predictive accuracy (77%) than all existing models, including DeepSol S2 (68%), DeepSol S3 (70%), and PaRSnIP (70%). There is a 16.2% and 12.7% difference in sensitivity between soluble and insoluble classes, for DeepSol S2 and DeepSol S3, respectively, where the insoluble class is predicted with higher accuracy. By contrast, the difference for the DSResSol (2) model is less than 6%, representing the outstanding capability of DSResSol (2) for identifying both soluble and insoluble classes, and thereby reducing prediction bias.

The DSResSol (1) model performs comparably to DeepSol S2 and DeepSol S3 models, and outperforms other models such as PaRSnIP and DeepSol S1. Performance of DSResSol (1) is competitive with DeepSol S2 and DeepSol S3; notably however, in contrast to DeepSol S2 and DeepSol S3, DSResSol (1) obtains a similar performance without additional biological features as complimentary information in the training process. The accuracy of DSResSol (1) is only 1% lower than DeepSol S2, and higher by at least 4% in accuracy and 7% in MCC than DeepSol S1, suggesting that our proposed model architecture can capture more meaningful information from the protein sequence than DeepSol S1, by using only protein sequence as input for the training process.

Figure 4 (A) and Figure 4 (B) show the Receiver Operating Characteristic (ROC) curve and the recall vs. precision curve for seven different solubility predictors, using the first independent test set *[30].* The area under curve (AUC) and area under precision recall curve (AUPR) for DSResSol (2) is 0.871 and 0.872, respectively, which are at least 2% higher than other models confirming that the DSResSol (2) model outperforms other state-of-the-art available predictors. Figure 4 (C) shows the accuracy of models in different probability threshold cutoffs. The highest accuracy for DSResSol (2) and DSResSol (1) is achieved at probability thresholds equal to 0.5. This achieved result is due to using a balanced training and testing set.

**Figure 4.**
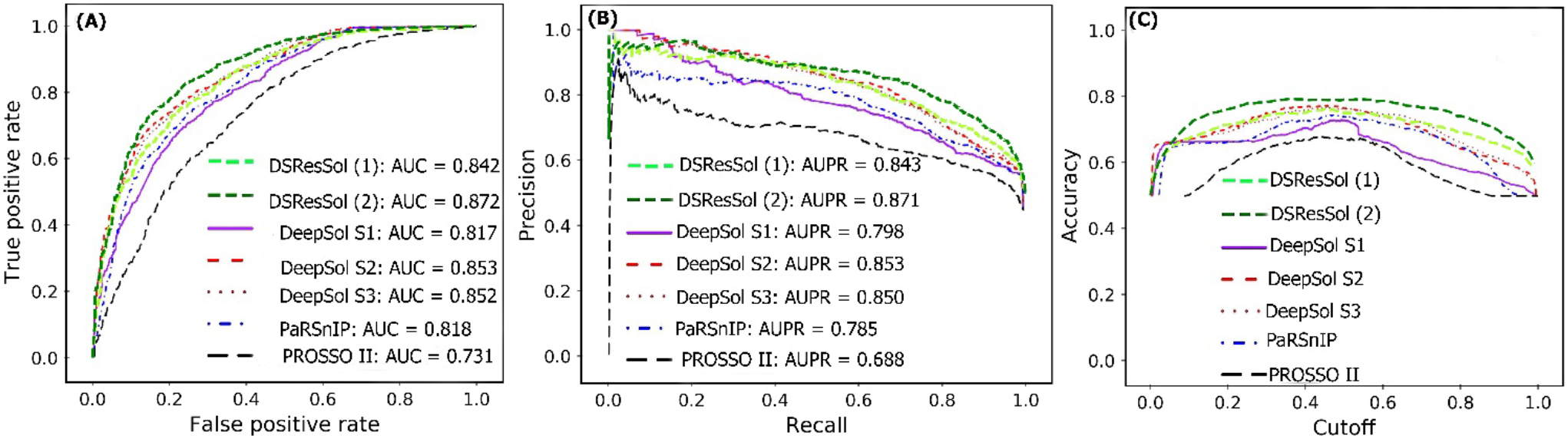
Comparison of the performance of DSResSol models with DeepSol [15] and PaRSnIP [13] models. (A) Receiver operating curve (ROC), (B) recall-precision curve, (C) accuracy-threshold cutoffs curve. The cutoff threshold discriminates between the soluble and the insoluble proteins. The curve for PROSSO II is obtained with permission from Bioinformatics [15].

For the second test set, (the newest test set to date), we evaluate the performance of both models and compare their performances with eight different available sequence-based tools. Table 2 represents the performance of the DSResSol models on the NESG test set. Evaluation metrics include both threshold-dependent metrics such as accuracy and MCC as well as threshold-independent metrics such as area under ROC curve value. We find that the accuracy and MCC of DSResSol (2) is at least 5% and 50% more than SoluProt tools, respectively. Also, DSResSol (2) achieves the highest AUC value (0.68) among other tested solubility predictors on the second independent test set. The DSResSol (1) model, using only protein sequences for training, achieves comparable results with other tools such as SoluProt [14] and PROSSO II [12]. The accuracy value for DSResSol (1) only is 2% lower than the best competitor (SoluProt). This demonstrates an outstanding performance of the DSResSol (1) model which does not take advantages of using additional biological features for training, confirming that DSResSol (1) indeed captures the most meaningful features from the protein sequence to distinguish soluble proteins from insoluble ones. Furthermore, the sensitivity of DSResSol (2) for soluble proteins is 73% which is significantly higher than SoluProt (at 13%). Figure 5 (A) and 5 (B) display the threshold-independent evaluation metrics to show the performance of our models in comparison to three different existing models, for the second independent test set (NESG) [27]. The area under ROC curve (AUC) and area under precision-accuracy curve (AUPR) are 0.683 and 0.678, respectively, which is at least 8% higher than the best existing competitor, SoluProt. In Figure 5 (C), we show the accuracy of the tested models at different solubility thresholds The highest accuracy (62%) for DSResSol models is obtained at the solubility threshold equal to 0.5.

**Figure 5.**
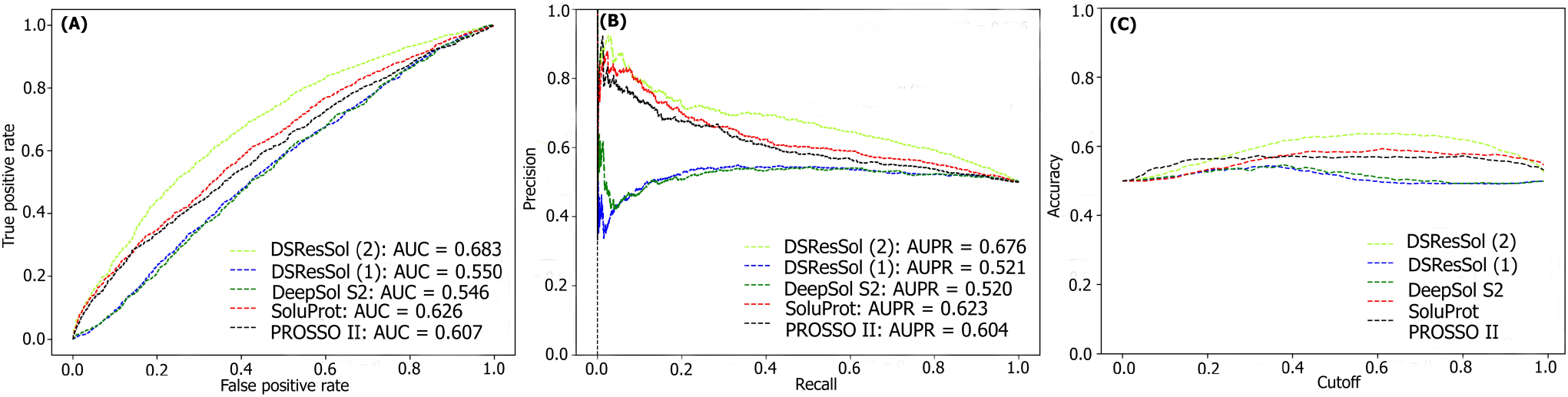
Comparison of the performance of DSResSol models with DeepSol**[15]**, PROSSO II [12], and SoluProt [14] models. (A) Receiver operating curve (ROC), (B) recall-precision curve, (C) accuracy-threshold cutoffs curve. The cutoff threshold discriminates between the soluble and the insoluble proteins.

We also consider the probability score distribution to evaluate DSResSol on both test sets. For the first test set, we consider the probability score distribution of DSResSol and close competitors (PaRSnIP [13] and DeepSol [15]) in violin plots (Figure 6) for both soluble and insoluble proteins. Distribution of scores for the four models shown in Figure 6 do not follow a normal distribution. For soluble and insoluble proteins, the score distribution plot shows that although DeepSol S2 like DSResSol (1) and DSResSol (2) gives more than a 99% level of confidence for the solubility prediction, the density of scores for DSResSol (2) in soluble proteins (values near score = 1) and for insoluble proteins (values near score = 0) is much higher than for DeepSol S2, confirming the better performance of DSResSol over DeepSol S2. In contrast to DSResSol and DeepSol S2, the score distribution for PaRSnIP model does not reach to score =1 for soluble and 0 for insoluble proteins, resulting in poor performance for PaRSnIP. To compare score distribution of DSResSol (1) and DeepSol S2, we can see that near the probability score cutoff = 0.5, DSResSol (1) has greater density scores than DeepSol S2, suggesting lower accuracy in comparison to DeepSol S2. We also compute the mean score for each model. For the insoluble class, the mean score of DSResSol (2) (0.12) is significantly lower than DeepSol S2 (0.26) and PaRSnIP (0.37). For the soluble class, the mean score for DSResSol (2) (0.81) is much higher than DeepSol S2 (0.62) and PaRSnIP (0.61), suggesting improved performance of DSResSol (2) over close competitors. In other words, a lower mean score value for the insoluble class and a higher mean score value for the soluble class represent a higher confidence in the model. Finally, we note that the density of scores in the DSResSol (1) model beyond a score of 0.5 is higher and lower for the insoluble and soluble classes, respectively, than DSResSol (2), suggesting lower accuracy (Figure 6). This result suggests that DSResSol (1) wrongly predicts more solubility values than DSResSol (2). This result can be understood considering that DSResSol (2) takes advantage of 85 additional biological features to establish a more accurate predictive model. For the second test set, we have done a similar analysis. We found that the DSResSol (2) outperforms both DeepSol S2 [15] and SoluProt [14] for the soluble class. However, for the insoluble class, DSResSol (2) has slightly inferior performance to DeepSol S2 for the second test set.

**Figure 6.**
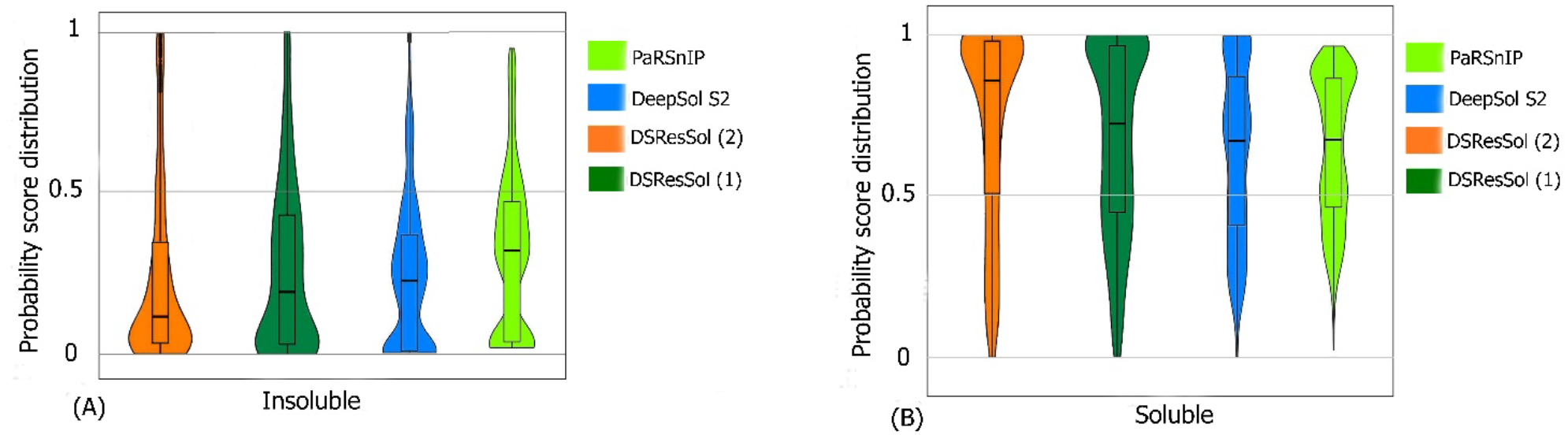
Violin plots represent the probability score distribution of DSResSol (1) and (2), DeepSol S2 [15], and PaRSnIP [13] for (A) insoluble and (B) soluble classes in the first test set [41].

For the second test set, we compare the probability score distribution with two competitors, SoluProt [14] and DeepSol [15]. Figure 7 shows this analysis for the second test set in two violin plots. The violin plot for insoluble class represents That the density of scores distribution for DeepSol S2 model is higher than DSResSol (2) and DSResSol (1) in probability score value equal to zero (probability score value = 0), indicating that the DeepSol S2 works better than DSResSol (2) for insoluble proteins prediction. Also, the density of scores distribution near the value = 0.5 for DSResSol is higher than DeepSol S2, confirming slightly better performance of DeepSol S2 on insoluble protein prediction in comparison to DSResSol (2). Furthermore, the mean of scores distribution of the SoluProt model is 0.63 for insoluble proteins, representing its relatively poor performance on insoluble proteins. However, based on Figure 7 B, DSResSol (2) outperforms DeepSol S2 and SoluProt for the soluble class. The mean of scores distribution on soluble proteins is 0.72 for DSResSol (2) while this value for DeepSol S2 and SoluProt is 0.42 and 0.66, respectively, confirming the DSResSol (2) is the best candidate tool for soluble proteins prediction. Furthermore, for soluble class, the density of scores distribution in DSResSol (2) is higher than both DeepSol S2 and SoluProt close to probability score value = 1, which further validate the outstanding performance of DSResSol (2) on soluble class.

**Figure 7.**
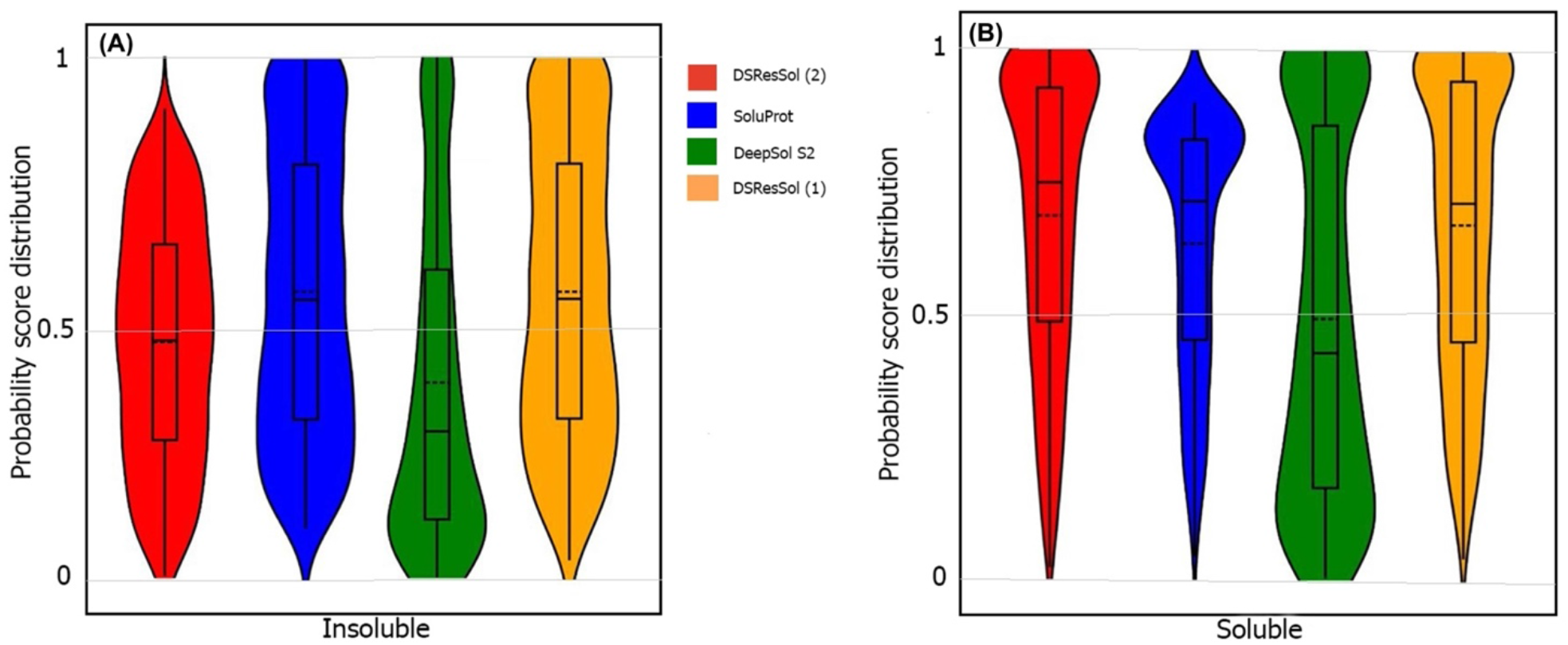
Violin plots represent the probability score distribution of DSResSol (1) and (2), DeepSol S2 [15], and SoluProt [14] for (A) insoluble and (B) soluble classes in NESG test set [27].

### 5.2 Effect of sequence length on solubility prediction

To illustrate the effect of protein sequence length on protein solubility prediction, we divide both test sets into five separate sets of different sequence length, in the range: {(0,100), (100, 200), (200, 300), (300, 400), (400, ∞)}. To evaluate how sequence length affects the solubility score, we show the score distribution for proteins predicted to be soluble and insoluble in five different sequence length ranges (Figure 8). True Positive (TP) and True Negative (TN) predictions correspond to the soluble and insoluble classes predicted correctly. False Positive (FP) and False Negative (FN) predictions correspond to the insoluble and soluble classes predicted incorrectly. Figure 8 (A) shows that the median decreases monotonically as sequence length increases, suggesting that longer sequence length results in reduced solubility. In other words, the shorter protein sequences are more soluble as proposed by Kramer et al. [42]. The median score for TP sets for five sequence length ranges are 0.93, 0.92, 0.90, 0.86, and 0.83, respectively. These values highlight the outstanding performance of DSResSol on the soluble class (a value of 1 corresponds to soluble protein). From Figure 8 (B), we observe that an increase in sequence length in the TN sets yields a decrease in the score distribution for insoluble proteins, suggesting that DSResSol can more easily predict insoluble proteins having longer than shorter sequences. The median of TN sets for five sequence length ranges is 0.21, 0.18, 0.16, 0.08, and 0.05, respectively, showing good performance of the DSResSol predictor (a value of 0 corresponds to insoluble protein). We further calculated the difference between median TN and FN (Median (FN) -Median (TN)) for the insoluble class as well as the difference between median of TP and FP (Median (TP) – Median (FP)) for the soluble class (Table 4). Proteins in the soluble class in the sequence length range (0, 100) and proteins in the insoluble class in the sequence length range (400<L<∞) have maximum values (0.29 for soluble and 0.34 for insoluble), confirming that the DSResSol model can predict proteins in these sequence length ranges with higher relative confidence than proteins with other sequence length ranges.

**Figure 8.**
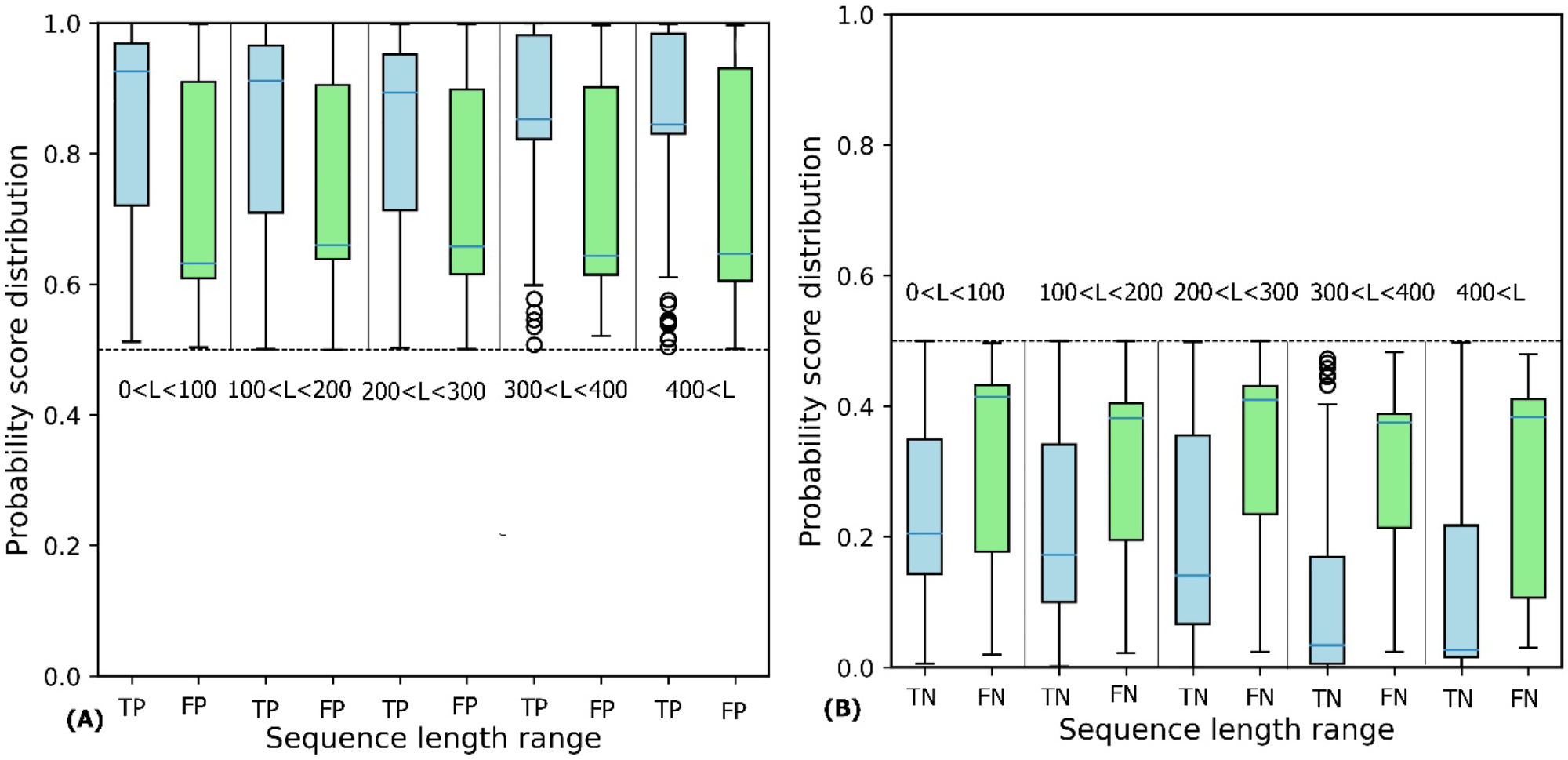
Probability score distributions for proteins predicted in both test sets to be (A) soluble and (B) insoluble for 5 different sequence length ranges: 0< L <100, 100< L <200, 200< L <300, 300<L<400, and 400< L<∞. TP = True Positive, FP = False Positive, TN = True Negative, and FN = False Negative. Blue horizontal line in box plot of each set shows the median of the score distribution for that set.

**Table 4.** Difference between True Positive and False Positive (TP, FP) as well as False Negative and True Negative (FN, TN) for different sequence length ranges. M = Median.

| Sequence length range | M(TP) – M(FP) | M(FN) – M(TN) |
| --- | --- | --- |
| (0,100) | 0.29 | 0.23 |
| (100,200) | 0.24 | 0.22 |
| (200,300) | 0.24 | 0.26 |
| (300,400) | 0.22 | 0.31 |
| (400,∞) | 0.22 | 0.34 |

### 5.3 Key amino acids, dipeptides, and tripeptides for protein solubility

To investigate the most important amino acids and di- and tripeptides contributing to protein solubility, we extract these directly from the DSResSol model. As discussed, nine initial CNNs in DSResSol are responsible for capturing amino acid k-mers for k = 1 to 9. The feature maps obtained from each initial CNN, having dimension 1200×32, are associated with amino acid k-mers for the corresponding protein sequence. To extract key amino acids associated with protein solubility, we need to obtain the feature vector, called activation vector, for each protein sequence. We extract these feature vectors for each protein sequence in our training set as follows. First, we pass the feature maps, which we receive from the CNN layer having filter size of 1, through a reshape layer to assign features maps with dimension 32×1200. Then, these feature vectors are fed to a Global Average Pooling layer to obtain the feature vectors of length 1200 for each protein sequence, which represents the activation vector for that protein sequence. Each value in the activation vector, called activation value, is associated with a corresponding amino acid within the original protein sequence. Hence, higher activation values suggest larger contribution to the classification results and protein solubility. We count the amino acids corresponding to the top 20 activation values for each protein sequence in the training dataset. The total number of each amino acid corresponding to the top 20 activation values for all protein sequences in the training dataset represent the importance of that amino acid in protein solubility classification. We apply the same process for feature maps obtained from initial CNN layers with filter size of 2 and 3 and count the total number of pairs and triplets corresponding to the top 20 activation value across all protein sequences to gain insight about the contribution of di- and tripeptides in protein solubility prediction. Figure 9 depicts the most important amino acids, dipeptides, and tripeptides contributing to protein solubility. From Figure 9 (A), we observe that glutamic acid, serine, aspartic acid, asparagine, histidine, and glutamine are key amino acids contributing to protein solubility. Glutamic acid, aspartic acid, and histidine are amino acid which have electrically charged side chains, while serine, asparagine, and glutamine have polar uncharged side chains. Interestingly, in one experimental study reported by Trevino et al., glutamine, glutamic acid, serine, and aspartic acid contribute most favorably to protein solubility [43]. Also, Figure 9 (B) and Figure 9 (C) show that two and three consecutive glutamine amino acids (EE and EEE) are the most important dipeptides and tripeptides contributing to protein solubility. These results are consistent with experimental data proposed by Islam et al. [44]. Additionally, polar residues and residues that have negatively charged side chains such as glutamic acid and aspartic acid are, in general, more likely to be solvent exposed than other residues [43], and can bind water better than other residues [45]. In another investigation, Chan et al. has demonstrated that positively charged amino acids like histidine are correlated with protein insolubility [46]. Also, Nguyen et al. found negatively charged fusion tags as another way to improve protein solubility [47], [48], consistent with our findings.

**Figure 9.**
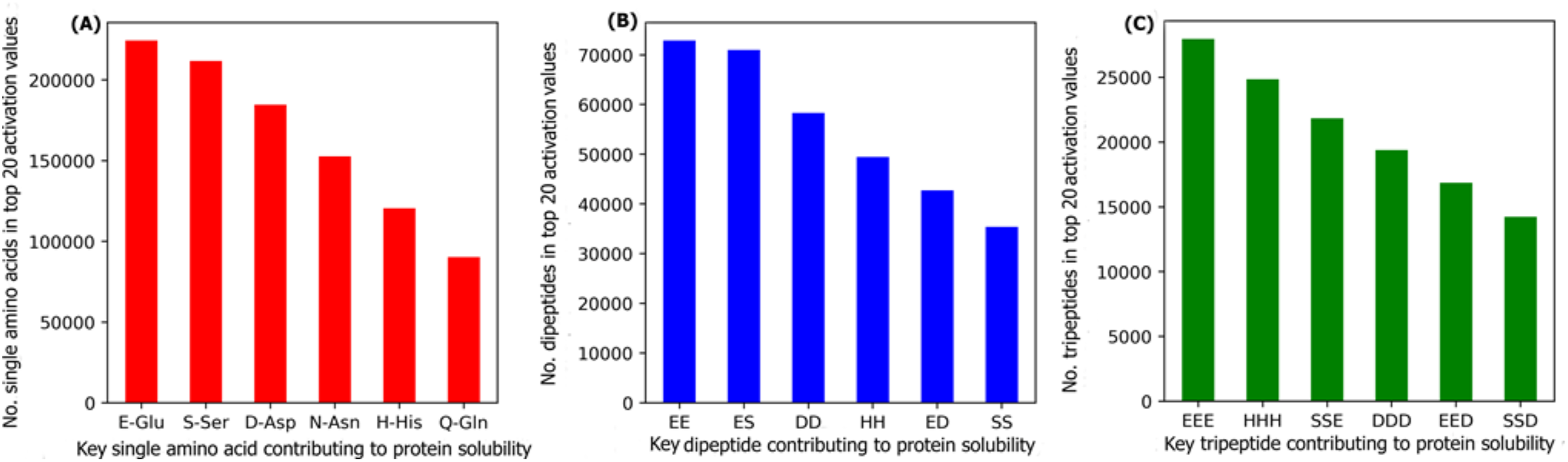
The number of (A) amino acids (B) dipeptides, and (C) tripeptides corresponding to the top 20 activation values, obtained from the initial CNNs in DSResSol model, across all protein sequences within the training dataset.

### 5.4 Effect of additional biological features on DSResSol performance

To evaluate the effect of each additional biological feature group on DSResSol performance, we consider each feature group independently in the DSResSol (1) model (SI Table 5 and SI Table 6). When we add only solvent accessibility related features to DSResSol (1), the accuracy of the model on the first test set increases from 0.751 to 0.782, and the accuracy of the model on the second test improves from 0.557 to 0.618. Adding secondary structure related features to DSResSol (1) improves the accuracy for first test set from 0.751 to 0.763 and for second test set from 0.557 to 0.582. We further analyze the fraction of exposed residues and secondary structure content for soluble and insoluble proteins in the training data. We identify that the soluble protein class has 61.2% helix and beta strand content. 68.7% of the residues are exposed residues with relative solvent accessibility cutoff higher than 65%. On the other hand, for the insoluble proteins in the training set, 81% of the secondary structure content is random coils. Further, 78% of residues are buried with relative solvent accessibility less than 35%, suggesting that the proteins having highly ordered structure and solvent exposed residues with larger relative solvent accessibility cutoffs, have a greater tendency to be soluble. By contrast, proteins with a higher degree of disordered secondary structure, such as random coil, and buried residues are predominantly insoluble. These results represent the influence on protein solubility propensity by solvent accessibility and secondary structure, and are supported by experimental data. Kramer et al. have previously demonstrated the correlation between solvent accessibility and secondary structure content with protein solubility. They proposed that soluble proteins have larger negatively-charged surface area, and thus amenable to bind water [42]. Also, Tan et al. identified the significant relationship between protein solubility and ordered secondary structure content such as helix and beta sheets [49]. They found large helix and beta sheet content within the most soluble proteins [49]. Thus, our results, which suggest that ordered secondary structures such as helix and beta sheets, as well as a larger fraction of solvent exposed residues with higher relative solvent accessibility cutoffs, contribute to protein solubility, correlate well with experimental findings.

### 5.5 Effect of sequence identity cutoff on DSResSol performance

To develop our input datasets, we removed the redundant protein sequences in our training set with sequence identity over 25%. Also, we eliminated the protein sequences in our training set having more than 15% sequence similarity with both test sets. To analyze the impact of identity cutoff on DSResSol performance, we consider different cutoffs to train DSResSol (SI Table 7 and SI Table 8). The results indicate that by changing the sequence identity cutoffs, the performance of DSResSol predictor improves to 75.1% for first test set and to 55.7% for second test set, suggesting that the optimal identity cutoff is 25% [15].

## 6. Discussion

In this study, we propose a novel sequence-based solubility predictor that uses a SE-ResNet neural network. In the first model, DSResSol (1), only raw protein sequences are used to distinguish soluble proteins from insoluble proteins. In the second model, DSResSol (2), to improve the performance of the first proposed model, 85 pre-extracted biological features including sequence as well as various structural features, are added. We observe that the performance of DSResSol (2) is superior to existing state-of-the-art tools when we evaluate the model performance on two different independent test sets. In particular, for first test set the accuracy of DSResSol (2) is at least 3% higher over the best performing model to date, DeepSol S2 [15]. For second test set, the accuracy of DSResSol is more than 5% higher than SoluProt [14], the top performing existing tool on this test set.

The main reason for the improved performance of the DSResSol predictor in comparison to other existing models originates from the SE-ResNet architecture. DeepSol used only some parallel CNNs to extract feature maps from the input protein sequence. In fact, the DeepSol model could only capture contextual features, and amino acid k-mers of different length and their local non-linear interactions. By contrast, our DSResSol model not only extracts amino acid k-mers and their local interactions but also captures long-range interactions between amino acid k-mers with different lengths. This is because DSResSol greatly benefits from the specific SE-ResNet architecture, including dilated CNNs. SE-ResNet blocks in DSResSol model are responsible for capturing frequently occurring amino acid k-mers where k = {1, 2,…9} and their local and global interactions. In fact, extracting these high order k-mers and their interactions gives valuable structural features such as protein folds [50], which are discriminative and crucial features as described in PaRSnIP [13]. In the SE-ResNet block, the dilated CNN efficiently extracts long-range interaction among k-mers, while preventing over-fitting using dropout on the weights, leading to good generalization performance. Also, SE-ResNet captures more information from the input feature maps related to amino acid k-mers because it not only reduces gradient vanishing owing to feature reusability, but also highlights the most important information from feature maps, which results in the capture of complex sequence-contact relationships while using many fewer parameters than other methods [29]. In addition, by adding 85 biological features to the DSResSol model the performance of the predictor is significantly improved. This suggests that these pre-extracted features are complementary to contextual features obtained from the SE-ResNet model.

In our analysis, we employ the DSResSol model to identify the relationship between sequence length and solubility propensity. For both insoluble and soluble classes, we observe monotonically decreasing score distributions when sequence length increases, suggesting that proteins with longer sequence have a higher tendency to be insoluble. Also, by analyzing the DSResSol model results, we find that the glutamine, serine, and aspartic acid are key amino acids that favorably contribute to protein solubility. Interestingly, this result is correlated with experimental studies reported by Islam et al. and Trevino et al. [43], [44]. Furthermore, we find that secondary structure and relative solvent accessibility features are determinative in protein solubility prediction. We demonstrate that soluble proteins include a large number of exposed residues at relative solvent accessibility cutoffs of more than 65%, and residues having ordered secondary structure content such as helix and beta sheets. On the other hand, for insoluble proteins, large number of residues are buried and disordered. These results are supported by experimental findings proposed by Kramer et al [42] and Tan et al [49].

## 7. Conclusion

In conclusion, by taking advantage of SE-Residual neural networks, we introduced the sequence-based solubility predictor, DSResSol (Dilated Squeeze excitation Residual network Solubility predictor), that outperforms all available bioinformatic tools for solubility prediction when the performance is assessed by different evaluation metrics such as accuracy and MCC. Also, in contrast to all existing models, DSResSol works well to identify both soluble and insoluble proteins (assessed by close sensitivity values for soluble and insoluble classes), suggesting that the superior accuracy of DSResSol originates from the great performance on both classes. Based on this observation, the model can be effectively used for soluble and insoluble identification of protein targets to avoid time-consuming trial-and-error approaches in experimental designs. Another interesting property of the DSResSol model is its outstanding performance in protein prioritization. Another reason for the model’s robustness originates not only from its novel deep learning architecture, but also from its comprehensive training dataset. In fact, we employ a novel training set that is cleaned from the noisy Target Track data via multiple steps for removing redundant protein sequences. We validate the model on two different independent test sets. In the future, the proposed architecture of DSResSol can be used for other protein feature predictions because this architecture can not only accurately extract but also boost the most meaningful features from the protein sequence. By incorporating these extracted features to other external features related to a specific property, other powerful predictors are currently in development. Furthermore, this model can be utilized uniquely for feature extraction from other computed properties. For example, rather than calculating the percentage of 3- and 8-state secondary structure content to obtain only 11 secondary structure related features as traditionally done, the DSResSol architecture can extract much more meaningful features from the complete secondary structure sequences. Therefore, the proposed architecture can be employed not only as an independent bioinformatics tool for other protein feature predictions but also as a feature extraction tool to prepare meaningful input features for other predictors.

