## Supplementary Information for "DSResSol: A sequence-based solubility predictor created with dilated squeeze excitation residual networks"


### 1. Dataset construction

SI Table 1. The number of sequences retained in each dataset construction step.

| Construction step | Training set | Soluble | Insoluble | Test set 1 | Soluble | Insoluble | Test set 2 | Soluble | Insoluble |
|---|---|---|---|---|---|---|---|---|---|
| Input | 129,593 | - | - | 2,001 | 1,000 | 1,001 | 9,703 | - | - |
| Pre-processing and solubility assignment | 109,648 | - | - | 2,001 | 1,000 | 1,001 | - | - | - |
| Redundancy removal | 87,969 | 40,905 | 14,064 | 2,001 | 1,000 | 1,001 | 9,423 | 5,718 | 3,705 |
| Removal of short sequences and sequences with unknown residues | 82,902 | 50,004 | 32,898 | 2,001 | 1,000 | 1,001 | 9,420 | 5,715 | 3,705 |
| Removal of transmembrane proteins | 76,274 | 45,603 | 30,671 | 2,001 | 1,000 | 1,001 | 8,769 | 5,421 | 3,348 |
| Removal of insoluble sequences with available PDB structure | 72,756 | 42,530 | 30,226 | 2,001 | 1,000 | 1,001 | 8,754 | 5,421 | 3,333 |
| Clustering to 25% identity | 49,369 | 26,422 | 22,947 | 2,001 | 1,000 | 1,001 | 3,945 | 2,078 | 1,867 |

| | | | | | | | | | |
|---|---|---|---|---|---|---|---|---|---|
| Overlap removal with test sets 15% identity | 46028 | 24920 | 21108 | 2,001 | 1,000 | 1,001 | 3,945 | 2,078 | 1,867 |
| Class and length balancing | **40,317** | 19,718 | 20,599 | **2,001** | 1,000 | 1,001 | **3,729** | 1,864 | 1,865 |

### 2. Hyperparameters in the model

SI Table 2. Tested hyperparameters in network layers and optimal values generated via the Grid Search Method [1] .

| **Layers** | **Number of tested units or filters** | **Optimal Value** | **Filter size** | **Parameters** | **Tested Values** | **Optimal Value** |
|---|---|---|---|---|---|---|
| Embedding layer | (50, 100, 150) | 50 | - | Epochs | 50 | 50 |
| Initial CNNs | (32, 64, 128, 256) | 32 | {1, 2, …, 9} | Learning rate | (0.005,0.008,0.01, 0.02) | 0.008 |
| Dilated CNN | (32,64,128,256) | 32 | 3 | Batch size | (32,64,128,256) | 64 |
| Bottlenecked CNN | (32,64,128,256) | 32 | 1 | Decay rate | (10e -7,10e-8, 10e-9) | 10e-7 |
| Final CNNs | (32,64,128,256) | 32 | {11, 13, 15} | Early stopping value | (3,4,5,6) | 5 |
| FC | (64,128,256,512) | 128 | - | - | - | - |
| MaxPooling | (2,3,5,7) | 3 | - | - | - | - |

### 3. Performance of 10 models obtained from 10-fold cross validation for DSResSol (1) and DSResSol (2)

The validation accuracy for each model is obtained from validation data. We utilize 10-fold cross validation for the training process. The training data is divided into 10 parts: 9 parts for training and 1 part for validation, alternatively in each cross-validation. Thus, in each cross-validation training, we use 10% of the training data as validation data and the remaining 90% of the data for the training process.

SI Table 3. Performance comparison for 10 models obtained from 10-fold cross-validation for DSResSol (1) models on both independent test sets. Red blocks are for the first test set and blue blocks are for the second test set. Validation accuracy is on validation data.

| Model | Training ACC | Valid ACC | ACC | Precision | Recall | F-1 score | ACC | Precision | Recall | F-1 score |
|---|---|---|---|---|---|---|---|---|---|---|
| 1 | 0.761 | 0.753 | 0.745 | 0.748 | 0.745 | 0.745 | 0.549 | 0.553 | 0.542 | 0.545 |
| 2 | 0.764 | 0.762 | 0.749 | 0.753 | 0.748 | 0.749 | 0.551 | 0.550 | 0.547 | 0.548 |
| 3 | 0.763 | 0.758 | 0.741 | 0.748 | 0.741 | 0.746 | 0.539 | 0.538 | 0.549 | 0.544 |
| 4 | 0.765 | 0.761 | 0.751 | 0.754 | 0.750 | 0.751 | 0.556 | 0.558 | 0.551 | 0.553 |
| 5 | 0.769 | 0.759 | 0.744 | 0.748 | 0.743 | 0.744 | 0.536 | 0.537 | 0.542 | 0.539 |
| 6 | 0.765 | 0.760 | 0.742 | 0.747 | 0.743 | 0.744 | 0.541 | 0.539 | 0.550 | 0.541 |
| 7 | 0.759 | 0.758 | 0.750 | 0.754 | 0.749 | 0.751 | 0.549 | 0.551 | 0.549 | 0.548 |
| 8 | 0.774 | 0.768 | 0.745 | 0.749 | 0.743 | 0.746 | 0.552 | 0.550 | 0.547 | 0.537 |
| 9 | 0.771 | 0.769 | 0.749 | 0.753 | 0.747 | 0.750 | 0.557 | 0.558 | 0.542 | 0.552 |
| 10 | 0.768 | 0.758 | 0.743 | 0.748 | 0.743 | 0.744 | 0.532 | 0.529 | 0.541 | 0.529 |

SI Table 4. Performance comparison for 10 models obtained from 10-fold cross-validation for DSResSol (2) models on both independent test sets. Red blocks are for the first test set and blue blocks are for the second test set. Validation accuracy is on validation data.

| Model | Training ACC | Valid ACC | ACC | Precision | Recall | F-1 score | ACC | Precision | Recall | F-1 score |
|---|---|---|---|---|---|---|---|---|---|---|
| 1 | 0.814 | 0.805 | 0.789 | 0.783 | 0.790 | 0.787 | 0.619 | 0.608 | 0.611 | 0.610 |
| 2 | 0.816 | 0.799 | 0.787 | 0.786 | 0.789 | 0.787 | 0.620 | 0.612 | 0.613 | 0.612 |
| 3 | 0.814 | 0.802 | 0.791 | 0.788 | 0.789 | 0.788 | 0.621 | 0.618 | 0.601 | 0.611 |
| 4 | 0.809 | 0.808 | 0.796 | 0.788 | 0.790 | 0.789 | 0.629 | 0.620 | 0.616 | 0.618 |
| 5 | 0.808 | 0.810 | 0.787 | 0.784 | 0.788 | 0.786 | 0.608 | 0.610 | 0.599 | 0.605 |
| 6 | 0.810 | 0.799 | 0.793 | 0.789 | 0.787 | 0.788 | 0.625 | 0.622 | 0.616 | 0.619 |
| 7 | 0.806 | 0.790 | 0.791 | 0.787 | 0.790 | 0.789 | 0.629 | 0.622 | 0.618 | 0.621 |
| 8 | 0.819 | 0.806 | 0.790 | 0.784 | 0.787 | 0.786 | 0.592 | 0.598 | 0.607 | 0.601 |
| 9 | 0.820 | 0.795 | 0.784 | 0.786 | 0.788 | 0.787 | 0.609 | 0.608 | 0.603 | 0.604 |
| 10 | 0.818 | 0.811 | 0.785 | 0.787 | 0.785 | 0.786 | 0.617 | 0.614 | 0.612 | 0.613 |

SI Table 5. Performance of the DSResSol model after adding each biological feature group to the DSResSol (1) model for the first test set. The accuracy of DSResSol (1) without biological features is 0.751.

| **Model** | **ACC DSResSol (1) after adding the additional biological features** | **ACC Improvement** |
|---|---|---|
| DSResSol (1) + Solvent accessibility related features | 0.787 | 3.7% |
| DSResSol (1) + Secondary structure related features | 0.762 | 1.1% |
| DSResSol (1) + order/disorder related features | 0.757 | 0.6% |
| DSResSol (1) + global sequence features | 0.756 | 0.5% |

SI Table 6. Performance of the DSResSol model after adding each biological feature group to the DSResSol (1) model for the second test set. The accuracy of DSResSol (1) without biological features is 0.557.

| **Model** | **ACC DSResSol (1) after adding the additional biological features** | **ACC Improvement** |
|---|---|---|
| DSResSol (1) + Solvent accessibility related features | 0.618 | 6.1% |
| DSResSol (1) + Secondary structure related features | 0.582 | 2.5% |
| DSResSol (1) + order/disorder related features | 0.564 | 0.7% |
| DSResSol (1) + global sequence features | 0.561 | 0.4% |

SI Table 7. Performance comparison for DSResSol (1) on the first independent test set for different cutoff sequence identity. Note: Best performing method in bold.

| **Model** | **ACC** | **MMC** | **Sensitivity (Soluble)** | **Sensitivity (Insoluble)** |
|---|---|---|---|---|
| DSResSol (1) Cutoff 25% | **0.751** | **0.508** | **0.691** | **0.813** |

| | | | | |
|---|---|---|---|---|
| DSResSol (1) Cutoff 15% | 0.744 | 0.491 | 0.686 | 0.805 |
| DSResSol (1) Cutoff 20% | 0.743 | 0.488 | 0.701 | 0.795 |
| DSResSol (1) Cutoff 30% | 0.744 | 0.492 | 0.688 | 0.801 |

SI Table 8. Performance comparison for DSResSol (1) on the second independent test set for different cutoff sequence identity. Note: Best performing method in bold.

| **Model** | **ACC** | **MCC** | **Sensitivity (Soluble)** | **Sensitivity (Insoluble)** |
|---|---|---|---|---|
| DSResSol (1) Cutoff 25% | **0.557** | **0.166** | 0.545 | **0.568** |
| DSResSol (1) Cutoff 15% | 0.553 | 0.157 | 0.542 | 0.567 |
| DSResSol (1) Cutoff 20% | 0.555 | 0.164 | **0.547** | 0.563 |
| DSResSol (1) Cutoff 30% | 0.552 | 0.159 | 0.541 | 0.567 |
